# A rare *IL2RA* haplotype identifies SNP rs61839660 as causal for autoimmunity

**DOI:** 10.1101/108126

**Authors:** Daniel B. Rainbow, Marcin Pekalski, Antony J. Cutler, Oliver Burren, Neil Walker, John A. Todd, Chris Wallace, Linda S. Wicker

## Abstract

*IL2RA* is associated with multiple autoimmune diseases including type 1 diabetes (T1D). Higher expression of *IL2RA* mRNA and its protein product CD25 in T lymphocytes is associated with a T1D-protective haplotype. Here we show that a rare variation of this haplotype that loses the protective allele at a single SNP, rs61839660, reduces *IL2RA* expression and T1D protection, identifying it as the causal factor in disease.

---

Common variants of the gene *IL2RA* encoding CD25, the alpha chain of the IL-2 receptor, are associated with multiple autoimmune diseases including type 1 diabetes^1^ (T1D) and multiple sclerosis^2,3^ (MS). The *IL2RA* genetic association is influenced by several distinct haplotypes, with the most protective T1D haplotype shared between T1D (odds ratio (OR) = 0.63) and MS (OR = 0.85) and termed “group A”^3^. The group A haplotype is defined by eight SNPs in high linkage disequilibrium (LD) (Supplementary Table 1)^3^, including SNPs rs12722495 and rs61839660 that correlate with higher CD25 expression on CD4^+^ memory T cells when the minor alleles are present^4–6^. One study demonstrated increased *IL2RA* mRNA from the A haplotype in the same cell type^4^, with the effect large enough to be seen as an eQTL in whole blood^7^, and the same SNP correlating with a *cis*-meQTL 250 bp distal of the *IL2RA* transcriptional start site^8^. Although SNPs in the group A haplotype have been identified as the most associated in genetic association studies of T1D ^1,9^and Crohn’s disease (rs61839660)^10^ as well as juvenile idiopathic arthritis (rs7909519)^11^, owing to the high degree of LD amongst the eight SNPs (minimum pairwise r^2^=0.65), the most associated SNP is not necessarily the causal variant^12^, and resolution of high LD between the most disease-associated variants in a region remains a major barrier to causal variant identification.

Epigenetic evidence suggests rs6183990 as the most likely causal variant as analysis of the ENCODE^13^ and Roadmap Epigenetic^14^ databases showed that it was the only A haplotype SNP that lies within conserved DNA sequence with potential enhancer activity in CD4^+^ T cells (Fig. 1a, Supplementary Fig. 1). The Regulome and Haploreg databases assigned rs61839660 as the most likely SNP from the eight SNP group A haplotype to alter transcriptional regulation of *IL2RA* (Supplementary Table 1). Promoter capture Hi-C data also indicate that rs61839660 is located in a region of DNA that interacts with the *IL2RA* promoter in CD4^+^ T cells and therefore provides a potential mechanism for how this intronic SNP could affect the transcriptional regulation of *IL2RA*^15^ (http://www.chicp.org).

**Figure 1:**
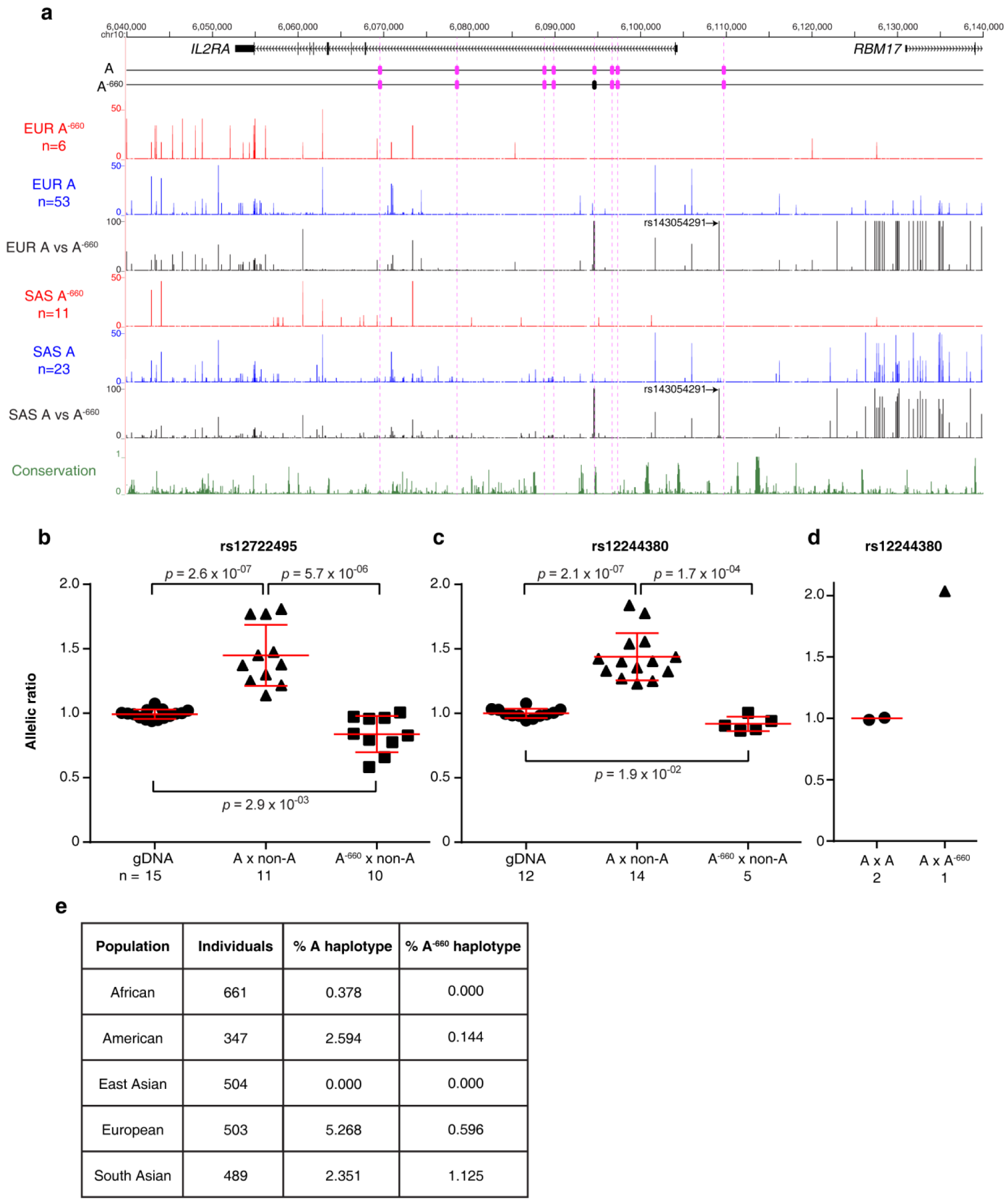
SNP rs61839660 determines allele-specific expression of *IL2RA*. (A) The location of the eight group A haplotype SNPs (pink ovals). In the A^−660^ haplotype the T allele of rs61839660 is replaced by C (black oval). Genotype similarity across 3,123 SNPs from donors carrying either the A or A^−660^ haplotype identified from the 1000 Genomes Project using the European (EUR) and South Asian (SAS) populations. The height of each line represents the minor allele frequency on the A and A^−660^ haplotypes for each SNP (see Methods). Conservation track displays multiple alignments of 100 vertebrate species and measurements of evolutionary conservation calculated by PhastCons and represent probabilities of negative selection. (B-D) ASE of *IL2RA* mRNA in CD4^+^ central memory T cells. The allelic ratio of *IL2RA* mRNA made by the chromosome carrying the A or A^−660^ haplotype compared to that carrying the non-A haplotype was measured using reporter SNP rs12722495 (B) located in intron 1 of *IL2RA*, and rs12244380 (C-D) located within the 3′ UTR of *IL2RA*. Red lines show mean and SD. P values were calculated using a Wilcoxon rank sum test. (E) Table showing the frequency of the A and A^−660^ haplotypes within the 1000 Genomes Project^18^.

We chose a novel approach to test whether SNP rs61839660 causally affects the expression of *IL2RA* mRNA, using cells from individuals with a rare *IL2RA* haplotype identified from the Cambridge BioResource, a large panel of volunteers recallable by specific genotypes. By phasing dense genotype information across the *IL2RA* region we identified a rare variation (0.6% frequency, Fig. 1e) of the T1D-protective group A haplotype in which all of the protective alleles are present except at rs61839660, where the protective T allele has been replaced with the susceptible C allele (called A^-660^, Fig. 1a, Supplementary Table 2). Comparison of the A and A^-660^ haplotypes using 1000 Genomes Project whole-genome sequence data revealed that the A and A^-660^ haplotypes share a remarkably similar genetic background (across 70 kb, chr10:6,055,500-6,126,500; Fig. 1a) suggesting the A and A^-660^ haplotypes have arisen together through an ancient bottleneck event. Through this many non-ancestral alleles (including the A haplotype SNPs) have been fixed, when compared to the susceptible haplotype that shows a greater proportion of the ancestral alleles present (Supplementary Fig. 2). One additional SNP (rs143054291) differed between the A and A^-660^ haplotypes across all individuals (Fig. 1a). As this A^-660^ haplotype-unique SNP is found on all donors having the A^-660^ haplotype from different world populations, it suggests a single gene conversion event^16^ created the A^-660^ haplotype and SNP rs143054291 arose at a very similar point in time. We infer that the gene conversion event introduced a short piece of DNA from a non-A haplotype replacing the DNA segment including rs61839660 as well as an in/del polymorphism 92 bp centromeric to the SNP (Supplementary Fig. 3). Based on this sequence comparison, we hypothesised that if rs61839660 does alter CD25 expression on CD4^+^ memory T cells^4–6^, we would expect A^-660^ donors to have CD25 expression levels comparable to the non-A haplotypes, which carry none of the group A T1D protective alleles^3^.

**Figure 2:**
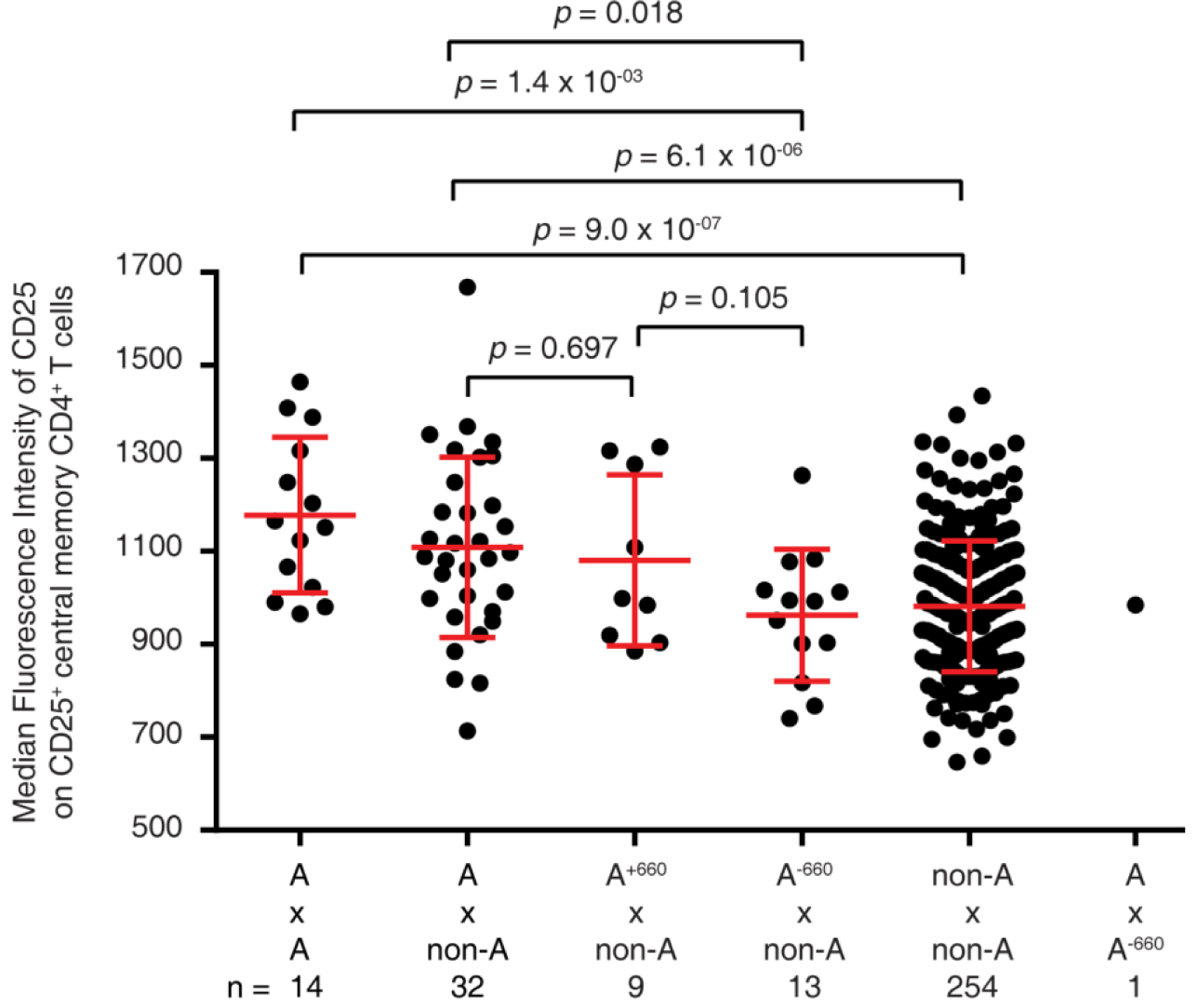
CD25 expression on CD4^+^ central memory T cells is determined by SNP rs61839660. The median fluorescence intensity of surface CD25 expression on CD4^+^ central memory T cells is displayed for individuals grouped by *IL2RA* diplotype. The gating strategy is shown in Supplementary Fig. 2. Non-A haplotypes are those not having any of the T1D-protective alleles at the eight group A SNPs. The red lines represent the mean with SD. P values were calculated using a Wilcoxon rank sum test.

The rarity of the A^−660^ haplotype (Fig. 1e) makes the study of A^−660^ homozygous individuals impractical even in the large panel of donors with ImmunoChip-based genotypes available through the Cambridge BioResource (n=2748). We therefore developed a high throughput method to measure allele-specific expression (ASE) to assess the relative amount of *IL2RA* mRNA produced from each chromosome within individuals heterozygous for the A or A^−660^ haplotypes and a non-A haplotype (A x non-A, and A^−660^ x non-A, respectively). The ASE method developed uses next-generation sequencing (NGS) and is based on a method we developed to perform targeted analysis of methylation^17^. To validate our method to measure ASE, we first reproduced the finding that the chromosome carrying the group A protective haplotype produces more *IL2RA* mRNA in CD4^+^ memory T cells compared to non-A haplotypes^4^ when measured by NGS in A x non-A donors using a reporter SNP in intron 1 (rs12722495, allelic ratio 1.44, *p*=2.6x10^−07^, Fig. 1b) or in the 3′ UTR (rs12244380, allelic ratio 1.44, *p*=2.1x10^−07^, Fig. 1c). Next we assessed the ASE in A^−660^ x non-A donors in CD4^+^ memory T cells and found significantly reduced allelic ratios (0.84 at rs12722495, *p*=5.7x10^−06^, Fig. 1b; 0.92 at rs12244380, *p*=1.7x10^−04^, Fig. 1c), a level close to that from the genomic DNA control (allelic ratio ~1 at rs12722495 and rs12244380 Fig. 1b, Fig. 1c).

From within the Cambridge BioResource, we identified one recallable donor out of 2,748 individuals heterozygous for the A^−660^ and A haplotypes. ASE was performed using mRNA from purified CD4^+^ central memory T cells using only the reporter SNP rs12244380 since the donor was homozygous at rs12722495. Compared to cells from two A-homozygous donors, which displayed nearly equal mRNA expression from both alleles, cells from the A^−660^ x A donor produced approximately twice the amount of *IL2RA* mRNA from the A allele as compared to the A^−660^ allele (Fig. 1d,c).

Examination of the surface CD25 expression on CD4^+^ memory T cells from 323 healthy donors replicated the finding that CD25 surface expression increases proportionally with the number of A haplotypes present^4^ (Fig. 2). A^−660^ x non-A donors had similar levels of CD25 expression as donors carrying only non-A haplotypes, and lower levels than donors heterozygous and homozygous for the A haplotype (*p*=0.018 and *p*=0.0014, respectively, Fig. 2). Nine donors carrying rare recombinant A haplotypes that all have T1D-protective alleles at SNPs rs61839660, rs12722496 and rs12722495 (A^+660^; Supplementary Table 3) but not at all five of the other group A SNPs had a similar surface CD25 expression (*p*=0.697) to A x non-A donors.

Having shown that increased CD25 expression on central memory CD4^+^ T cells as well as increased *IL2RA* mRNA levels determined by the A haplotype were not attributes of the A^-660^ haplotype, we next tested the A^-660^ haplotype for T1D disease association. Despite the rarity of A^-660^ haplotypes, we observed lower protection from T1D for the A^-660^ compared to the A haplotype (OR = 1.37, CI=1.015-1.840, *p*=0.04, Supplementary Fig. 5).

Our study demonstrates the power of using quantitative measurements of allele-specific mRNA production from rare haplotypes present in the human population as a method to resolve the high LD between the most disease-associated variants in order to identify the causal variant. The A^-660^ haplotype is reminiscent of a natural CRISPR experiment, with the advantage that accessible cell types can be purified and analysed *ex vivo* in a physiological context. The fact that both the A and A^-660^ haplotypes have been maintained in some human populations but not others (Fig. 1e) suggests that the functional consequences of distinct *IL2RA* regulation by these two haplotypes can be advantageous or deleterious depending on the environment. The identification of the A^-660^ haplotype and targeted NGS-based ASE analysis facilitates the future study of rs61839660’s influence on *IL2RA* regulation in all primary cell types that express CD25 constitutively or following activation. The loss of protection from T1D by the A^-660^ haplotype as compared to the A haplotype (Supplementary Fig. 5) should be examined in other autoimmune diseases where the A haplotype at *IL2RA* is associated with protection or susceptibility^3,10,11^. Overall our findings highlight the value of dense genotyping and whole genome sequencing of recallable donors in bioresources to identify rare recombination or gene conversions events that may influence gene expression in a physiological context.

## Acknowledgements

This work was supported by Wellcome Trust Grants 096388 and 107212, JDRF Grants 9-2011-253 and 5-SRA-2015-130-A-N, the National Institute for Health Research Cambridge Biomedical Research Centre (BRC). CW is supported by the Wellcome Trust (107881) and the UK Medical Research Council (MC_UP_1302/5). We gratefully acknowledge the participation of all NIHR Cambridge BioResource volunteers. We thank the Cambridge BioResource staff for their help with volunteer recruitment. We thank members of the Cambridge BioResource SAB and Management Committee for their support of our study and the National Institute for Health Research Cambridge Biomedical Research Centre for funding. Access to Cambridge BioResource volunteers and to their data and samples are governed by the Cambridge BioResource SAB. Documents describing access arrangements and contact details are available at http://www.cambridgebioresource.org.uk. We thank Graeme Clark, Howard Martin, Fay Rodger and Ruth Littleboy for running the Illumina MiSeq in the Stratified Medicine Core Laboratory NGS hub, Department of Medical Genetics, Cambridge University, Addenbrooke’s Hospital, Cambridge. This research was supported by the Cambridge NIHR BRC Cell Phenotyping Hub. In particular, we wish to thank Anna Petrunkina Harrison, Simon McCallum, Christopher Bowman, Natalia Savinykh, Esther Perez and Jelena Markovic Djuric for their advice and support in cell sorting. We also thank Helen Stevens, Pamela Clarke, Gillian Coleman, Sarah Dawson, Jennifer Denesha, Simon Duley, Meeta Maisuria-Armer and Trupti Mistry for acquisition and preparation of samples.

### Author contributions

DBR, MP, AC and OB performed the practical work and analysed the data. CW performed statistical analyses of T1D genetic disease risk. DBR and LSW drafted the manuscript, and all authors contributed to the editing of the manuscript. LSW, JAT and CW supervised the work.

### Competing financial interests

The authors declare no competing financial interests.

